# Identification and prioritisation of causal variants in human genetic disorders from exome or whole genome sequencing data

**DOI:** 10.1101/209882

**Authors:** Nagarajan Paramasivam, Martin Granzow, Christina Evers, Katrin Hinderhofer, Stefan Wiemann, Claus R. Bartram, Roland Eils, Matthias Schlesner

**Author notes:** Shared senior authorship (Roland Eils, Matthias Schlesner).

## Abstract

With genome sequencing entering the clinics as diagnostic tool to study genetic disorders, there is an increasing need for bioinformatics solutions that enable precise causal variant identification in a timely manner.

**Background:** Workflows for the identification of candidate disease-causing variants perform usually the following tasks: i) identification of variants; ii) filtering of variants to remove polymorphisms and technical artifacts; and iii) prioritization of the remaining variants to provide a small set of candidates for further analysis.

**Methods:** Here, we present a pipeline designed to identify variants and prioritize the variants and genes from trio sequencing or pedigree-based sequencing data into different tiers.

**Results:** We show how this pipeline was applied in a study of patients with neurodevelopmental disorders of unknown cause, where it helped to identify the causal variants in more than 35% of the cases.

**Conclusions:** Classification and prioritization of variants into different tiers helps to select a small set of variants for downstream analysis.

## Introduction

Next generation sequencing (NGS) has proven to be a powerful technique to identify causal genes in rare genetic disorders [1,2]. The decline in sequencing cost has enabled the use of NGS for diagnostics in the broader clinical setting [3–5]. However, the lower cost of data generation resulted in a daunting task of managing, analyzing and interpreting large data sets [6]. The bioinformatics tools and pipelines need to be constantly improved to keep pace and to enable a speedy analysis of the NGS data.

There are more than 3 million variants present in individual genomes compared to the human reference genome [7]. For clinical sequencing projects this list needs to be reduced to a manageable number of candidate variants for downstream analysis. One strategy to effectively reduce the number of candidate variants is trio sequencing, meaning that the healthy parents are sequenced along with their affected children.

Generally, analysis workflows for trio- or pedigree-based analysis share three basic steps [8]. In step one, the raw sequence reads are mapped to the reference genome for each sample individually and variants are called for all samples together by identifying the differences to the reference genome and determining the genotype per sample. In step two, technical artifacts and variants which are common in the population are removed as these are highly unlikely to be the cause of rare genetic diseases. Even after this first filtering step plenty of variants remain as potential candidates, and prioritization of the variants in the candidate list using pedigree information and various variant annotations is required [8].

Here, we present a complete workflow for candidate variant identification and prioritization to analyze whole exome sequencing (WES) and whole genome sequencing (WGS) data generated from trios or from larger pedigrees.

## Material and Methods

We developed a workflow that performs the read alignment, variant calling, annotation, filtering and prioritization steps. A particular focus is put on various filtering and annotation steps to prioritize variants depending on the assumed inheritance model for further analysis.

### Read alignment

Raw sequencing data were mapped to the 1000 genome reference genome (hs37d5) [7] using BWA [9] aln version 0.6.2 with standard parameters except setting ‘-q 20’. The resulting SAM files were sorted, converted to BAM format and indexed using SAMtools-0.1.19 [10]. Multiple lanes per sample were merged and duplicate reads marked using Picard [11] with the following parameter settings, ‘picard-1.61 MarkDuplicates VALIDATION_STRINGENCY=SILENT REMOVE_DUPLICATES=FALSE ASSUME_SORTED=TRUE MAX_RECORDS_IN_RAM=12500000 CREATE_INDEX=TRUE CREATE_MD5_FILE=TRUE’.

### Variant calling and annotation

Single nucleotide variants (SNVs) and small indels (1-20 bps) were jointly called from all the samples in a family using Platypus [12] with following parameter settings, ‘Platypus_0.8.1.py callVariants nCPU=10 genIndels=1 genSNPs=1 minFlank=0-bamFiles=$List_of_Bam_Files’ --refFile=hs37d5.fa --output=$Output_VCF’. Gene and transcript definitions from Gencode v19 [13] were added using ANNOVAR [14] and the minor allele frequency (MAF) information from 1000 genome Phase III and Exome Aggregation Consortium (ExAC) [15] were added. In addition, allele frequencies from 328 WES and 177 WGS inhouse samples (referred to as local controls) were added to the variants.

### Variant filtering

Variants that passed all the Platypus internal filters were considered further. Frequent variants were removed based on MAF threshold of 0.1% from ExAC and the 1000 genome phase III database. To remove technical artifacts specific to our pipeline, variants that were present in the local controls above the threshold of 2% were considered as artifacts and were removed.

In the trio sequencing setting variant information from the patient and the healthy parents was combined to efficiently reduce the number of candidates. Only variants fulfilling an inheritance model for a disease were considered further. For an autosomal dominant (AD) disease, only *de novo* variants, i.e. variants which are present in the patient but not in any of the parents were considered as candidates. In case of autosomal recessive (AR) inheritance of consanguineous parents, we expected for the genotype to be homozygous in the patients and heterozygous in both parents. Hemizygous variants had to be heterozygous in only one of the parents and homozygous in the patient. Candidates for X-linked (XL) variants had to be heterozygous in the mother and hemizygous in the male patient. Finally, to identify candidates for compound heterozygous inheritance of AR diseases, SNVs and indels were combined to find two different heterozygous mutations in the patient in the same gene that were inherited one from each of the parents.

Variants were further checked for their genotype quality in all the samples in the family and low genotype quality variants (Phred score <20) were filtered out. The remaining variants were prioritized to select a list of candidate causal variants for further downstream analysis.

### Variant prioritization

Both variants and genes were prioritized by different measures and classified into different tiers, from which the final candidates were selected.

Variants were prioritized based on their effects on protein function, which were predicted from various conservation scores. To this end, various annotations from dbNSFP [16] including the GERP score [17] and CADD scores [18] were added to the variants. The GERP score [17] measures the evolutionary conservation of the sequences across species; a position with a score greater than two is considered as a highly conserved nucleotide and its disturbance is likely to have a high functional impact. CADD scores [18] integrate various annotations including sequence conservation scores and ENCODE project functional annotations to measure the deleteriousness of the variants. A CADD score of 13 in Phred scale was used as threshold, which means that prioritized variants are considered to be in the top 5% of the deleterious variants in the human genome.

Genes were prioritized based on their intolerance towards functional mutations. Intolerance missense z scores or pLI from ExAC were added if the variant was a missense or loss of function (LoF, includes stop gain/loss, splice acceptor/donor or frameshift indels), respectively. An increasing positive z-score indicates increasing intolerance to a missense mutation and we consider a z-score above +2 as an outlier intolerant gene, and a pLI score > 0.9 is considered intolerant to a LoF variant.

Finally, variants were categorized into two tiers which each contained three levels. Level 0 of both tiers contained the whole variant set before prioritization. In Tier 1 only LoF variants were moved to level 1. LoF variants with CADD score above the threshold were moved further into level 2. Finally, variants in level 2 which affect genes with pLI score above the threshold were moved into level 3. The missense variants in level 0 were moved into level 1 of Tier 2 and further prioritized into different levels of Tier 2, accordingly. Here, instead of the ExAC pLI score the ExAC missense Z-score is used to prioritize variants into level 3. In the downstream analysis, initially only the variants in level 3 of both tiers were considered. Only if there were no candidates found in level 3, variants in lower levels were examined.

In addition, these prioritized variants were further classified using the guidelines providing by the American College of Medical Genetics and the Association for Molecular Pathology [19] by medical geneticists during reporting.

**Figure 1:**
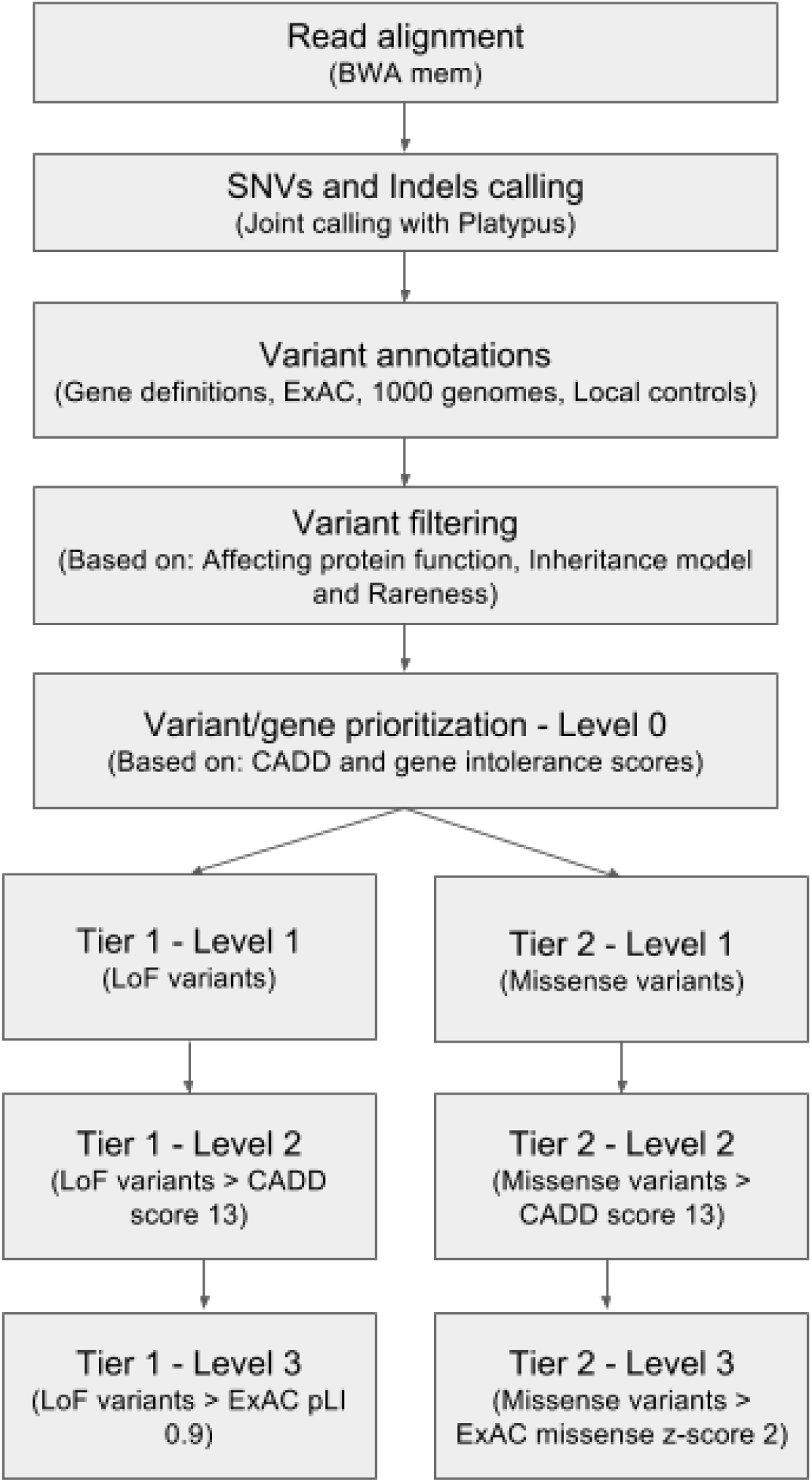
Germline variant analysis pipeline for rare genetic disorders

## Results

Recently we published results from our clinical exome initiative [4], where we analyzed exome data from 60 families with undiagnosed neurodevelopmental disorders (NDD), neurometabolic disorders (NMD) and dystonias and overall found causal variants in 35% of the families. In this current manuscript, we will use the results from 39 families in the NDD cohort to show how the germline analysis pipeline effectively filtered and prioritized variants identified from WES data in a trio sequencing setting.

Among the 127 WES samples from 39 families, on average 201,687 SNVs and 38,186 indels were present with a minimum coverage of 10X. Among them, 152,641 SNVs and 14,912 indels were present in ExAC or 1000 genome phase III with an MAF above 0.1%, or in our Local controls with a frequency above 2%, and were discarded. On average there were 6,368 SNVs and 451 indels remaining which were used for further analysis.

In the next step, all variants outside of exonic regions (+/− two base pairs to account for splice sites) as defined by the Gencode v19 gene model were removed. The remaining variants were classified on how they affect protein function and variants that cause missense and LoF mutations were selected further. On average there were 528 SNVs and 29 indels which are functional and rare/private remaining per sample.

For all 39 families of the NDD cohort, trio sequencing was performed. In some families in addition to the patient and parents, the affected or unaffected siblings were also sequenced, which enables more efficient variant filtering for recessive and X-linked diseases. For analysis, we considered such cases as separate trio families and only in the final steps of the pipeline the results were merged and then reported family-wise. The pedigree information was used to classify the variants into de novo, homozygous, hemizygous and heterozygous variants (Figure 2). The heterozygous SNVs and indels were further combined to find the compound heterozygous variants present in these families. At the end of the variant filtering steps, an average of 4 de novo, 5 homozygous, 1 hemizygous and 2 pairs of compound heterozygous small variants, respectively, remain per family. These variants were further prioritized into different tiers to select the best candidate causal variants/genes for further analysis and confirmation.

In Tier 1, 69 LoF variants were found in level 1 from all the samples, on average 1 LoF variant per sample, and 49 of those variants have CADD score greater than 13 were in level 2, and 12 variants were in genes that are intolerant to a new LoF variants in level 3 (Figure 3). In Tier 2, there are 529 total missense variants and 296 of them have CADD score above 13, among them 67 variants are in the genes with ExAC mis_z score greater than 2 (Figure 4).

Using this approach, the causal variants could be identified in 15 in the NDD cohort (table 3, [4]), The inheritance pattern was AD in 6 of these families, AR in 7 families, and XL in 2 families. Five of the AD causal variants and 2 XL causal variants were found in level 3 of Tier 1 or 2, where the variants and genes satisfied all the thresholds in the prioritization step. Among the 7 AR families 8 AR variants; including the IFT140 AR variant were found only in one case in the family 4, only one variant was in the top level of tier 2. The other AR causal variants had high variant deleteriousness scores, but the genes were predicted to be tolerant to new functional mutations. We have to note that the intolerance scores are not developed for AR inheritance models, so in general, for the AR disease inheritance, the gene intolerance score based prioritization level have to be relaxed to find the causal variants.

**Figure 2:**
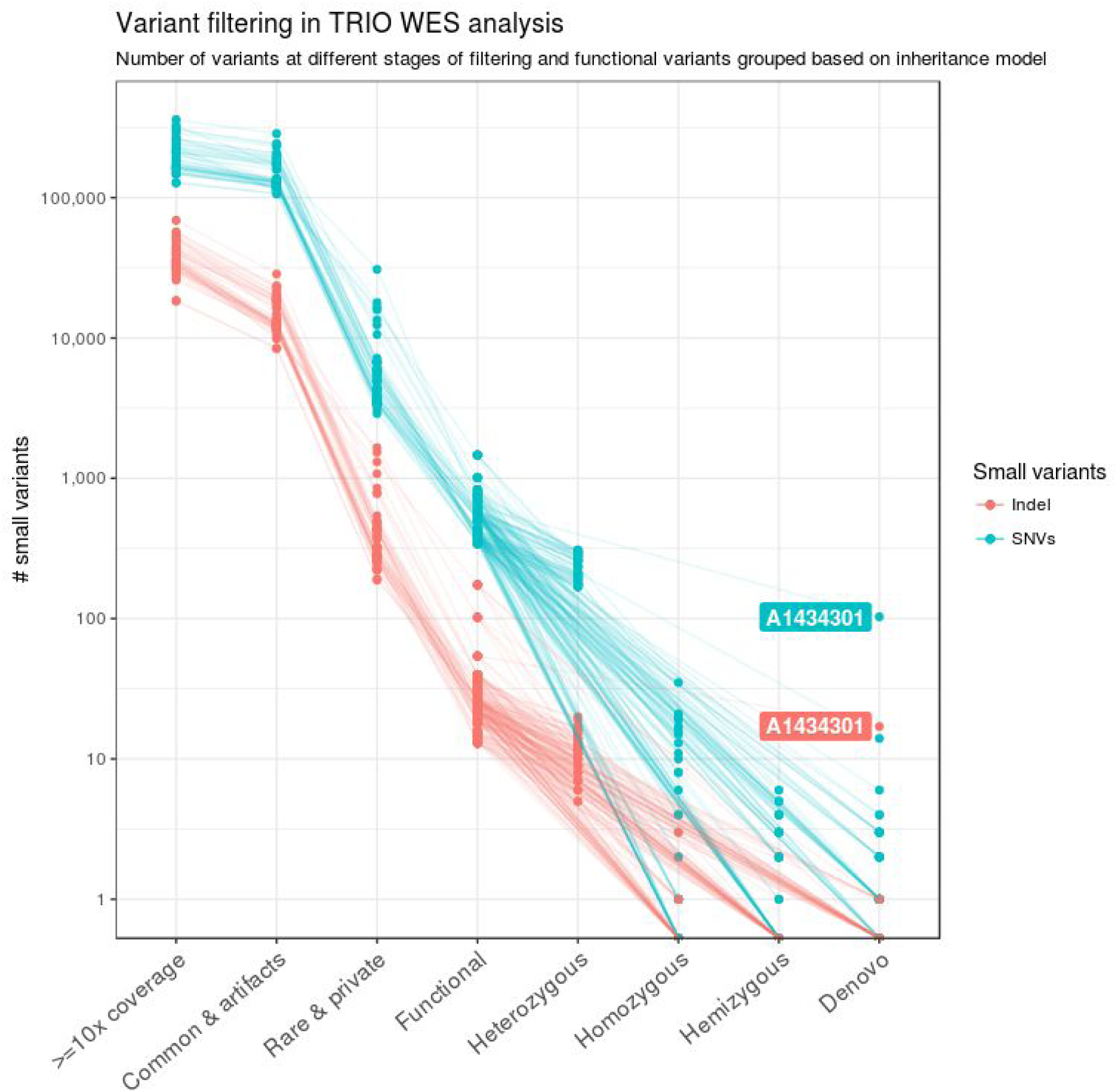
Variant filtering in TRIO WES analysis. The number of candidate variants after the different filtering steps is shown for the 39 families from the NDD cohort. After the basic quality filters, functional variants are identified and patient and parents data are combined. Based on the pedigree information the functional variants are further classified into different genetic models. The sample A1434301 has a high number of de novo variants due to poor DNA quality.

**Figure 3:**
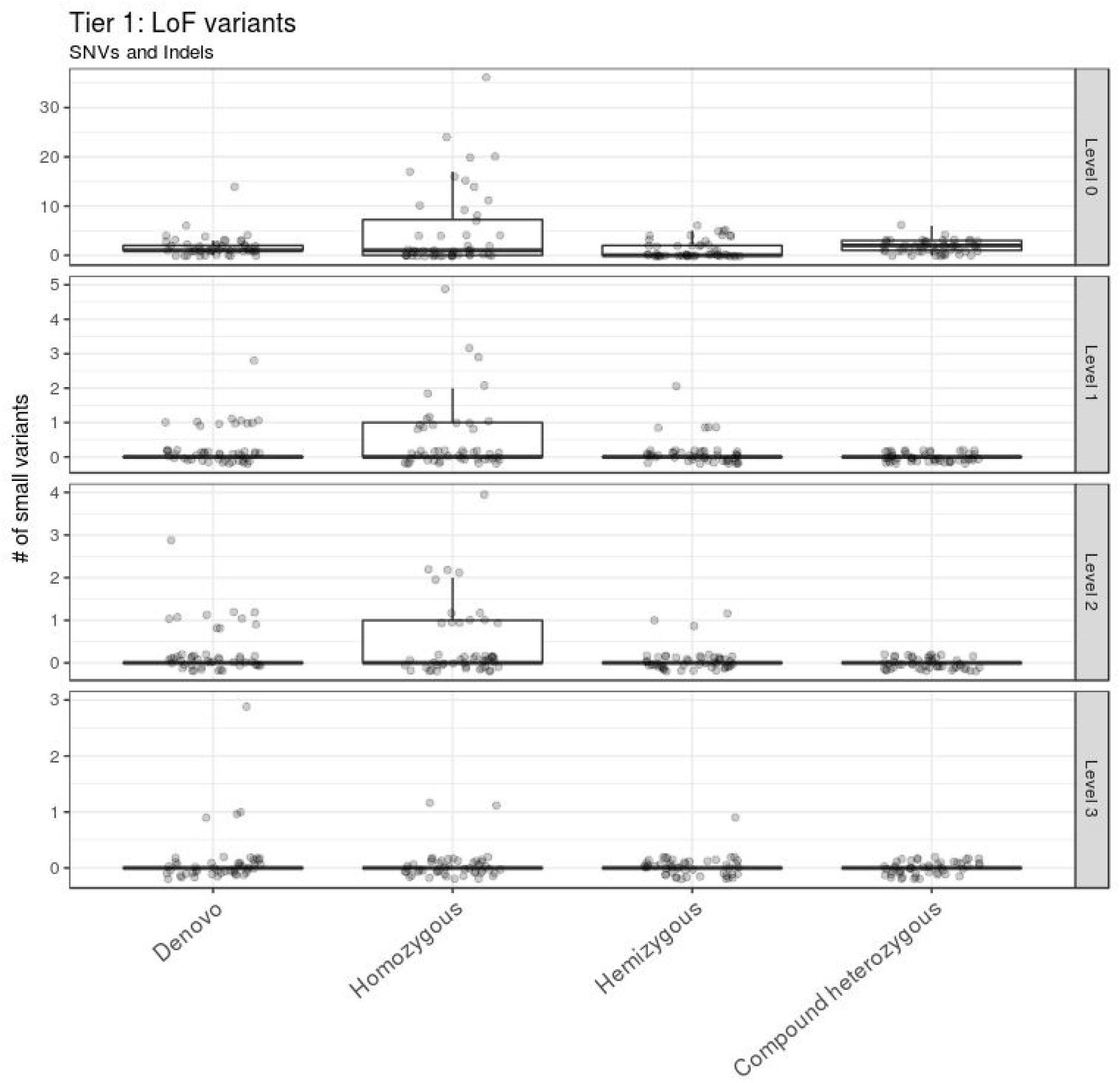
LoF variant prioritization. The LoF variants are classified into different levels in Tier 1. LoF variants with CADD score > 13 and ExAC pLI score > 0.9 are in level 3. They constitute the first set of candidate variants to be considered for further downstream analysis. The outlier A1434301 is removed from the visualization.

**Figure 4:**
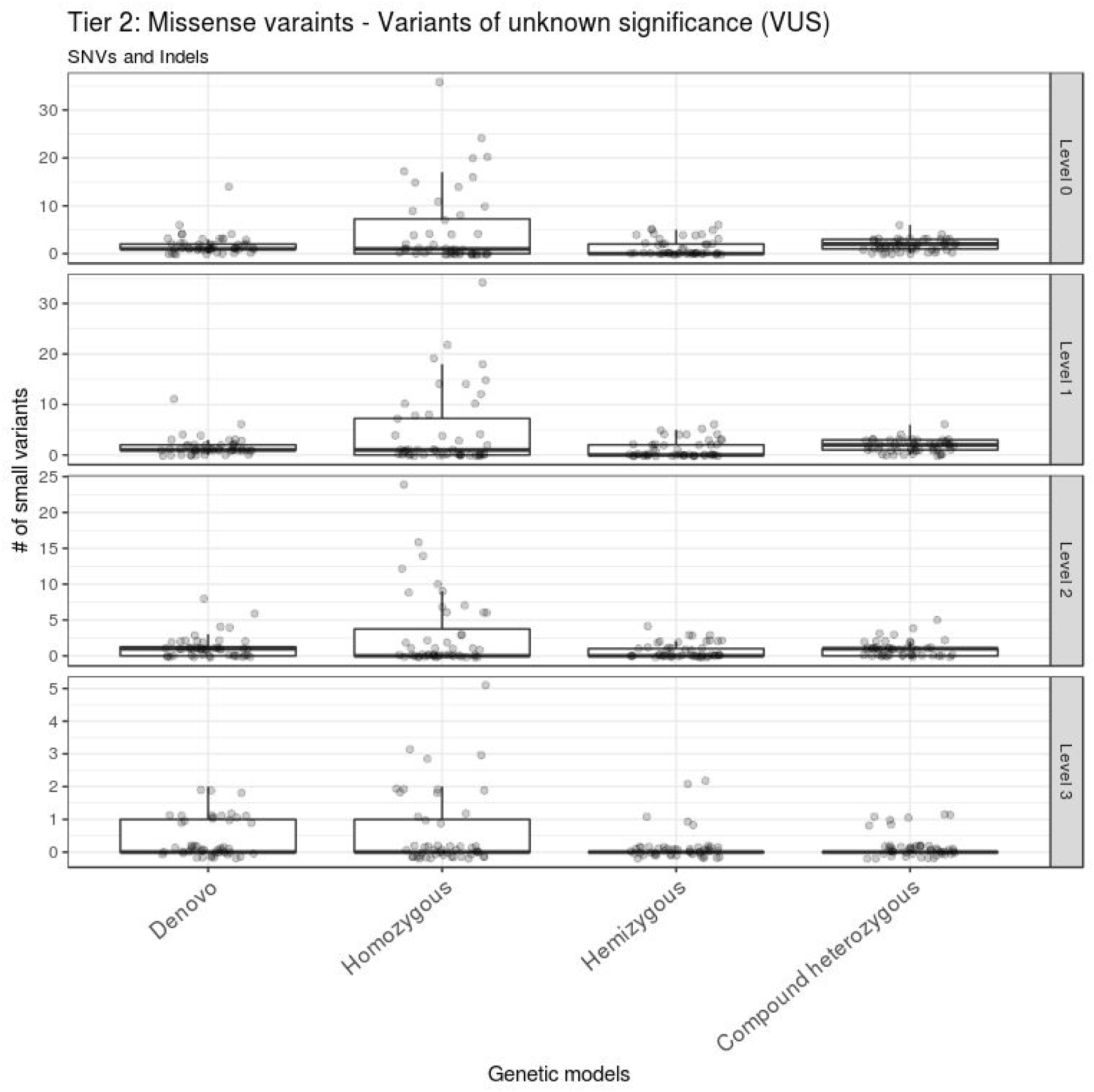
Missense variants prioritization. Missense variants are classified into different levels of Tier 2. Only missense variants with CADD score > 13 and ExAC missense z-score > 2 are in level 3. These variants constitute the second set of variants to be considered after the level 3 variants in Tier 1. The outlier A1434301 is removed from the visualization.

## Discussion

Next-generation sequencing offers the possibility to identify causal variants of genetic diseases without prior hypothesis about the affected gene. The rapidly decreasing costs for NGS allow now the application of whole exome or even whole genome sequencing as routine diagnostic tool in clinical settings. However, such hypothesis-free sequencing results in long lists of candidate variants. As manual variant filtering and candidate selection based on background knowledge is prohibitive in large studies and routine clinical application, there is a high need for bioinformatics workflows to extract short lists of prioritized candidates. Our germline analysis pipeline outlined here achieves this by making use of trio sequencing or larger pedigree sequencing data. Without the pedigree information, many variants will remain and it is much harder to pick a candidate from the prioritized gene list.

Generally, workflows for identification of disease-relevant variants rely on prefiltering steps to remove common polymorphisms and technical artifacts, usually using publicly available and private databases. Of note, the majority of the samples in the public databases like ExAC and 1000 genome are from the Central European Population (CEP). The majority of the samples analyzed in our study were also from CEP so the combined MAF from all populations was used. However, it is recommended to use population-specific MAF if the samples are from other populations than CEP. It is also recommended to aggregate AF from the samples sequenced on the same platform and processed with the same pipeline to remove technical artifacts arising specifically on the sequencing platform and pipeline used.

After the filtering steps, predictions about the functional impact of variants are used to narrow down the set of candidates. Many deleteriousness prediction tools are based on sequence conservation of the affected position [20]. However, individual tools are often not in a good agreement, so that it is recommended to use a consensus vote of multiple tools or to use meta-tools that combine the results of multiple individual prediction algorithms [8] [16]. For these reasons, our workflow relies on the CADD score [18], which integrates multiple levels of information including conservation and functional data, for variant prioritization.

A complementary information to the deleteriousness of the variant is the tolerance of the affected gene to new functional mutations. The intolerance or constraint score of a particular gene is calculated from the deviation of the observed number of functional mutations in a gene in large populations from the expected number which is based on the total amount of variation in this gene [21]. However, also the gene intolerance score can be misleading, as it does for example predict major cancer predisposition genes to be tolerant to new functional mutations [20]. Since there are subregions of the genes which are much more intolerant to functional mutations than other regions, a region-based or exon-based intolerance score could achieve a higher resolution and hence enable better predictions. Furthermore, as noted earlier in the results section, the gene intolerance score performs poorly for AR variants. So, a new gene intolerance score calculated solely from the homozygous variants in the gene should be used to prioritize variants in the AR inheritance model.

Current studies to identify causal variants in genetic diseases, focus usually only on exonic, functional variants and investigate only SNVs and small indels. In these studies, for up to 60% of the investigated cases no causal variants could be identified [3,4,22]. A possible explanation might be that in these cases the causal variants are focal copy number variants (CNVs) or structural variants (SVs), and might also be interested to look into +/− 10 bases around the splice sites, which cannot comprehensively be identified from WES data and thus require WGS for full exploration. Alternatively, causal variants might affect non protein-coding regions of the genome. While these variants can be identified from WGS data, their interpretation and functional effect prediction is much more challenging than for coding variants. Multiple efforts have been made recently to produce more accurate tools to predict effect of non-coding variants in Mendelian diseases [23,24], so that in the future the inclusion of non-coding variants into prioritization workflows should be possible.

## Conclusion

The rapid decrease in sequencing costs opened the door for broad application of genome sequencing in research as well as in clinical settings. The resulting massive sequence data production made data analysis and interpretation a daunting task in clinical genomics. We have developed an automated variant identification and prioritization pipeline for genetic disorders. With the classification of variants and genes into different tiers, the pipeline makes it easy to focus on a small set of candidate variants for downstream analysis. The pipeline will continuously be updated by adding new better accurate tools and scores to improve its performance and will eventually also allow to include SVs, CNVs, and non-coding variants.

## Acknowledgments

We thank and the Genomics and Proteomics Core Facility at the German Cancer Research Center for their excellent technical support and expertise. Authors like to acknowledge the data management group (DMG) in the Division of Theoretical Bioinformatics for managing the sequence data.

## Author Contributions

NP and MS developed the strategies and NP developed the pipeline. MS, CRB and RE provided the guidance and support. MG, CE and KH provided the samples and SW provided the sequencing data. NP, MS and RE wrote the manuscript. All the authors have read and contributed to the manuscript.

## Funding Source

This work was supported by the BMBF-funded Heidelberg Center for Human Bioinformatics (HD-HuB) within the German Network for Bioinformatics Infrastructure (de.NBI) (#031A537C).

## Competing Interests

Authors have no competing interests.

## Reference

1. Ng SB, Buckingham KJ, Lee C, Bigham AW, Tabor HK, Dent KM, et al. Exome sequencing identifies the cause of a mendelian disorder. Nat Genet. 2010 Jan;42(1):30–5.

2. Bamshad MJ, Ng SB, Bigham AW, Tabor HK, Emond MJ, Nickerson DA, et al. Exome sequencing as a tool for Mendelian disease gene discovery. Nat Rev Genet. 2011 Sep 27;12(11):745–55.

3. Lee H, Deignan JL, Dorrani N, Strom SP, Kantarci S, Quintero-Rivera F, et al. Clinical exome sequencing for genetic identification of rare Mendelian disorders. JAMA. 2014 Nov 12;312(18):1880–7.

4. Evers C, Staufner C, Granzow M, Paramasivam N, Hinderhofer K, Kaufmann L, et al. Impact of clinical exomes in neurodevelopmental and neurometabolic disorders. Mol Genet Metab. 2017 Aug;121(4):297–307.

5. Biesecker LG, Green RC. Diagnostic clinical genome and exome sequencing. N Engl J Med. 2014 Jun 19;370(25):2418–25.

6. Lelieveld SH, Veltman JA, Gilissen C. Novel bioinformatic developments for exome sequencing. Hum Genet. 2016 Jun 1;135(6):603–14.

7. 1000 Genomes Project Consortium, Auton A, Brooks LD, Durbin RM, Garrison EP, Kang HM, et al. A global reference for human genetic variation. Nature. 2015 Oct 1;526(7571):68–74.

8. Pabinger S, Dander A, Fischer M, Snajder R, Sperk M, Efremova M, et al. A survey of tools for variant analysis of next-generation genome sequencing data. Brief Bioinform. 2014 Mar 1;15(2):256–78.

9. Li H, Durbin R. Fast and accurate short read alignment with Burrows-Wheeler transform. Bioinformatics. 2009;25(14):1754–60.

10. Li H, Handsaker B, Wysoker A, Fennell T, Ruan J, Homer N, et al. The Sequence Alignment/Map format and SAMtools. Bioinformatics. 2009 Aug 15;25(16):2078–9.

11. Picard Tools - By Broad Institute [Internet]. [cited 2017 Aug 28]. Available from: http://broadinstitute.github.io/picard/index.html

12. Rimmer A, Phan H, Mathieson I, Iqbal Z, Twigg SRF, WGS500 Consortium, et al. Integrating mapping-, assembly- and haplotype-based approaches for calling variants in clinical sequencing applications. Nat Genet. 2014 Aug;46(8):912–8.

13. Harrow J, Frankish A, Gonzalez JM, Tapanari E, Diekhans M, Kokocinski F, et al. GENCODE: the reference human genome annotation for The ENCODE Project. Genome Res. 2012 Sep;22(9):1760–74.

14. Wang K, Li M, Hakonarson H. ANNOVAR: functional annotation of genetic variants from high-throughput sequencing data. Nucleic Acids Res. 2010;38(16):e164–e164.

15. Lek M, Karczewski KJ, Minikel EV, Samocha KE, Banks E, Fennell T, et al. Analysis of protein-coding genetic variation in 60,706 humans. Nature. 2016 Aug 18;536(7616):285–91.

16. Liu X, Jian X, Boerwinkle E. dbNSFP v2.0: A Database of Human Non-synonymous SNVs and Their Functional Predictions and Annotations. Hum Mutat. 2013;34(9):E2393–402.

17. Cooper GM, Goode DL, Ng SB, Sidow A, Bamshad MJ, Shendure J, et al. Single-nucleotide evolutionary constraint scores highlight disease-causing mutations. Nat Methods. 2010 Apr;7(4):250–1.

18. Kircher M, Witten DM, Jain P, O’Roak BJ, Cooper GM, Shendure J. A general framework for estimating the relative pathogenicity of human genetic variants. Nat Genet. 2014 Mar;46(3):310–5.

19. Richards S, Aziz N, Bale S, Bick D, Das S, Gastier-Foster J, et al. Standards and guidelines for the interpretation of sequence variants: a joint consensus recommendation of the American College of Medical Genetics and Genomics and the Association for Molecular Pathology. Genet Med. 2015 May;17(5):405–24.

20. Eilbeck K, Quinlan A, Yandell M. Settling the score: variant prioritization and Mendelian disease. Nat Rev Genet. 2017 Oct;18(10):599–612.

21. Petrovski S, Wang Q, Heinzen EL, Allen AS, Goldstein DB. Genic intolerance to functional variation and the interpretation of personal genomes. PLoS Genet. 2013 Aug 22;9(8):e1003709.

22. Retterer K, Juusola J, Cho MT, Vitazka P, Millan F, Gibellini F, et al. Clinical application of whole-exome sequencing across clinical indications. Genet Med. 2016 Jul;18(7):696–704.

23. Huang Y-F, Gulko B, Siepel A. Fast, scalable prediction of deleterious noncoding variants from functional and population genomic data. Nat Genet. 2017 Apr;49(4):618–24.

24. Smedley D, Schubach M, Jacobsen JOB, Köhler S, Zemojtel T, Spielmann M, et al. A Whole-Genome Analysis Framework for Effective Identification of Pathogenic Regulatory Variants in Mendelian Disease. Am J Hum Genet. 2016 Sep 1;99(3):595–606.

